# The soluble glutathione transferase superfamily: Role of Mu class in Triclabendazole sulphoxide challenge in *Fasciola hepatica*

**DOI:** 10.1101/2020.07.22.213892

**Authors:** Rebekah B. Stuart, Suzanne Zwaanswijk, Neil D. MacKintosh, Boontarikaan Witikornkul, Mark Prescott, Peter M. Brophy, Russell M. Morphew

**Author notes:** Corresponding author: Dr R Morphew. Institute of Biological, Environmental and Rural Sciences (IBERS) Aberystwyth University, Aberystwyth, Wales. SY23 3DA.

## Abstract

*Fasciola hepatica* (liver fluke), a significant threat to food security, causes global economic loss for the livestock production industry and is re-emerging as a food borne disease of humans. In the absence of vaccines the commonly used method of treatment control is by anthelmintics; with only Triclabendazole (TCBZ) currently effective against all stages of *F. hepatica* in livestock and humans. There is widespread resistance to TCBZ and detoxification by flukes might contribute to the mechanism. However, there is limited Phase I capacity in adult parasitic helminths and the major Phase II detoxification system in adults is the soluble Glutathione transferases (GST) superfamily. Previous global proteomic studies have shown that the levels of Mu class GST from pooled *F. hepatica* parasites respond under TCBZ-Sulphoxide (TCBZ-SO), the likely active metabolite, challenge during *in vitro* culture ex-host. We have extended this finding by using a sub-proteomic lead approach to measure the change in the total soluble GST profile (GST-ome) of individual TCBZ susceptible *F. hepatica* on TCBZ-SO-exposure *in vitro* culture. TCBZ-SO exposure demonstrated a FhGST-Mu29 and FhGST-Mu26 response following affinity purification using both GSH and S-hexyl GSH affinity resins. Furthermore, a low affinity Mu class GST (FhGST-Mu5) has been identified and recombinantly expressed and represents a novel low affinity mu class GST. Low affinity GST isoforms within the GST-ome was not limited to FhGST-Mu5 with second likely low affinity sigma class GST (FhGST-S2) uncovered through genome analysis. This study represents the most complete *Fasciola* GST-ome generated to date and has supported the sub proteomic analysis on individual adult flukes.

## Introduction

Fasciolosis, caused by the trematode liver flukes *Fasciola hepatica* and *F. gigantica*, is a foodborne zoonotic affecting grazing animals and humans worldwide (Andrews 1999). Liver fluke cause economic losses of over US$3 billion worldwide per annum to livestock via a decrease in production of milk, meat and wool, susceptibility to other infections, condemnation of livers, and mortality (Boray 1997). There are no commercial vaccines as yet available with Triclabendazole (TCBZ) currently the most commonly used fasciolicide due to activity against both adults and juvenile stage fluke (Brennan, Fairweather et al. 2007). TCBZ is absorbed in the rumen and passes through the blood to the liver where it is rapidly oxidised to the likely main active metabolites; Triclabendazole-sulphoxide (TCBZ-SO) (Alvarez, Solana et al. 2005) and Triclabendazole sulphone (TCBZ-SO_2_) (Alvarez, Solana et al. 2005; Alvarez, Moreno et al. 2009). Unfortunately, TCBZ resistant liver fluke are wide spread, with resistance first encountered in Australia; but it is now evident in Western Europe (Brennan, Fairweather et al. 2007) including the UK (Thomas, Coles et al. 2000).

At present our understanding of the mode of action and detoxification of TCBZ is fragmented and mechanisms underpinning resistance may need to be resolved in order to measure early TCBZ resistance in populations and thus preserve efficacy (Brennan, Fairweather et al. 2007). To this end, the glutathione transferase (GST) superfamily have been identified as the major Phase II detoxification system present in all parasitic helminths. GSTs have been implicated in both drug metabolism and resistance in other groups of organisms e.g. insects and human tumours (Hayes and Pulford 1995). Eight cytosolic GST classes have been identified across kingdoms; namely Alpha, Mu, Pi, Sigma, Theta, Kappa, Zeta, and Omega (Cvilink, Lamka et al. 2009). In *F. hepatica* GSTs belonging to four classes have been revealed by biochemistry and bioinformatics; Omega (ω), Mu (μ), Sigma (σ) and Zeta (ζ) (Chemale, Morphew et al. 2006; Morphew, Eccleston et al. 2012). Chemale *et al*. (2010) further reported that Mu class GST levels vary, with Mu class GST-1 reduced in abundance while Mu class GST-26 increased in TCBZ resistant and susceptible *F. hepatica* under TCBZ sulphoxide (TCBZ-SO) exposure. In addition, Scarcella *et al*. (2012) identified that fluke resistant to TCBZ expressed significantly higher levels of GST activity compared to susceptible flukes. Furthermore, an amino acid mutation in Mu Class GST-26 has been linked to a TCBZ resistant liver fluke strain (Fernandez, Estein et al. 2015). However, to date, there has not been a robust sub-proteomic study that compared the expression of GST isotypes in individual liver fluke under TCBZ-SO stress. Thus, we purified GSTs from the cytosol of single adult flukes using a combination of Glutathione (GSH) and S-Hexyl-GSH agarose, resolved GST isotypes by 2-DE and identified individual GSTs by MS/MS with the support of genomic and transcriptomic databases. As a consequence we have identified a novel Mu, Sigma and Omega class GST designated FhGST-Mu5, FhGST-S2 and FhGST-O2 respectively. FhGST-Mu5 has been cloned and expressed in a recombinant and active form and characterised.

## Materials and Methods

### *In vitro* TCBZ culture

Individual liver fluke from natural infections were collected and exposed to TCBZ-SO as described previously (Morphew, MacKintosh et al. 2014). In brief, live adult *F. hepatica* were collected from a local abattoir (Randall Parker Foods, Llanidloes, Wales, UK) and washed in PBS at 37° C. Fluke were washed for 1 h with PBS replacement every 15 mins. Post-washes, replicates of 10 adult, sized matched, worms were placed into fluke Dulbecco’s Modified Eagle Medium (DMEM) culture media containing 15 mM HEPES, 61 mM glucose, 2.2 mM Calcium acetate, 2.7 mM Magnesium sulphate, 1 µM serotonin and gentamycin (5 µg/ml) as previously described (Morphew, Wright et al. 2011). Flukes were maintained in culture at 37°C for 2 h (including transport to the laboratory) to establish a baseline protein expression profile. Upon completion of the initial 2 h incubation, culture media was replaced and supplemented with TCBZ-SO (LGC Standards, UK) at 50 µg/ml (Lethal dose) or 15 µg/ml (Sub-lethal dose) in DMSO (final conc. 0.1% v/v). For control samples only DMSO was added to a final volume of 0.1% v/v. Fluke cultures were then allowed to incubate at 37°C for a 6 h time period after which the media was refreshed, with DMSO and TCBZ-SO as required. Fluke cultures were incubated at 37°C for a further 6 h. A final refreshment of culture media was conducted and fluke cultures incubated for an additional 12 h at 37°C. Upon completion of the culture fluke were removed from the media and snap frozen individually in liquid N_2_. All samples were stored at -80°C until required.

### GST Assay and Purification

Individual adult *F. hepatica* were homogenised in an all-glass homogeniser on ice using 2 ml of lysis buffer containing 20 mM Potassium Phosphate pH 7.4, 0.1% v/v Triton-X 100 and EDTA-free protease inhibitors (Roche, Complete-Mini, EDTA-free). Samples were centrifuged at 100,000 x *g* for 45 minutes at 4°C to obtain the supernatant, the cytosolic fraction. Protein levels were quantified by the method of Bradford (1976). GST enzymatic specific activity was determined according to the conditions outlined by Habig *et al*. (1974) described previously (LaCourse, Perally et al. 2012) and stored at -80°C until needed. Specific activity data was log_10_ transformed prior to statistical comparison carried out by a two way ANOVA.

Cytosolic proteins were applied to a GSH-agarose (Sigma-Aldrich) or an S-hexyl-GSH-agarose (Sigma-Aldrich) affinity matrix and purified at 4° C according to the manufacturer’s instructions and as described previously (Morphew, Eccleston et al. 2012). Eluted proteins were concentrated using 10-kDa molecular weight cut off filters (Amicon Ultra, Millipore) and washed with ddH_2_O. All samples were quantified again by the method of Bradford (Sigma-Aldrich).

### Protein Preparation and 2-DE

IPG strips (7 cm, linear pH 3-10) were rehydrated with 100 μl of buffer (containing 8 M Urea, 2% w/v CHAPS, 33 mM DTT, 0.5% ampholytes pH range 3-10) plus 25 μl of sample protein and ddH_2_O to load 20 µg of GSH or Hexyl-GSH affinity bound proteins. Samples were in-gel rehydrated for 16 hrs and isoelectrically focused on 7 cm pH 3-10 IPG strips to 10,000 Vh on a Protean® IEF Cell (BioRad). After focusing, strips were then equilibrated for 15 min in reducing equilibration buffer (30% v/v glycerol, 6 M urea, 1% DTT) followed by 15 min in alkylating equilibration buffer (30% v/v glycerol, 6 M urea, 4% iodoacetamide). IPG strips were run upon SDS PAGE (12.5% acrylamide) using the Protean® II 2-D Cell (BioRad). Gels were then Coomassie blue stained (Phastgel Blue R, Amersham, Biosciences), and imaged on a GS 800 calibrated densitometer (BioRad). Quantitative differences between 2-DE protein spots were analysed using Progenesis PG220, software version 200 (Nonlinear Dynamics Ltd.), using 5 biological replicates. Spots were automatically detected on gels and manually edited. Normalisation of spots was calculated using total spot volume multiplied by the total volume (Moxon, LaCourse et al. 2010). All gel images were warped using manual matching before average gels (5 gels were used to make the average gels) for each treatment group were produced. Unmatched protein spots were also detected on appropriate gel comparisons. Two-fold differences between protein spots with a p<0.05 were considered significant when average gels were compared.

### Western Blotting

Following 2-DE, resolved proteins were transferred to nitrocellulose membranes. The nitrocellulose membrane was soaked in ddH_2_O for 1 minute. The gel, membrane, filter paper and porous pads were equilibrated in 1X Western Blot Transfer Buffer (NuPAGE Transfer Buffer, Life Technologies) for 20 min.

Proteins were transferred at 40 V for 2 h in 1 X Western blot transfer buffer (50 ml NuPage transfer Buffer, 850 ml ddH_2_O, 100 ml Methanol). To ensure proteins were transferred, the membrane was removed and stained with Amido black staining solution (0.1% w/v Amido black, 10% v/v Acetic Acid, 25% v/v isopropanol) for 1 min to detect the success of the transfer. The membrane was then washed with ddH_2_O. The membrane was then placed in Amido black de-stain (25% v/v isopropanol and 10% v/v acetic acid) for 30 mins. The membrane was imaged using the GS-800 calibrated densitometer (BioRad). Amido black stain was removed with several washes of Tris buffered saline, 1% v/v Tween 20 (TTBS).

The nitrocellulose membrane was blocked in blocking buffer (TTBS + 5% milk powder) overnight. The membrane was then washed with TTBS and then incubated with the primary antibody for 1-2 h. A 1:5,000 dilution and a 1:30,000 antibody dilution in blocking buffer was used for anti-Mu and anti-Sigma GST respectively. After incubation with the primary antibody the membrane was washed in TBS for 10 min. The membrane was washed twice more before incubating with the secondary antibody (Anti-goat IgG raised in rabbits for the Mu and Anti-rabbit IgG raised in goats for the Sigma) for 1-2 hours at a 1:30000 dilution in blocking buffer. The membrane was then washed 3 times in Tris buffered saline (TBS). Interacting spots or bands were detected using the 5-Bromo-4-chloro-3-indolyl phosphate (BCIP) in conjugation with Nitro blue tetrazolium (NBT), according to manufactures instructions. To develop, a 1:2 solution of BCIP:NBT in substrate buffer consisting of 0.1 M tris, 100 mM NaCl and 5 mM magnesium chloride, adjusted to pH 9.5. To cease the over development, membranes were rinsed in ddH_2_O. Blots were then scanned with a GS-800 calibrated densitometer (BioRad) and were imaged using Quantity One Version 4.6 software (BioRad, U.K.).

### Protein Identification

Protein spots were manually excised from the gels and in-gel digested with trypsin according to the method of Chemale *et al*. (2006). Tandem mass spectrometry (MSMS) were performed according to the method described by Moxon *et al*. (2010). Briefly, selected peptides from peptide digests were loaded onto a gold coated nanovial (Waters, UK), and sprayed at 800 – 900 V at atmospheric pressure and fragmented by collision induced dissociation using agron as the collision gas. Mass Lynx v 3.5 (Waters, UK) ProteinLynx was used to process the fragmentation spectra. Each fragmented spectrum was individually processed as follows; each spectrum was combined and smoothed twice using the SavitzkyGolay method ± 3 channels, background subtraction (polynominal order 15 and 10% below the curve). Each spectrum was exported and spectra common to each 2-DE spot were merged into a single MASCOT generic format (.mgf) file using the online peak list conversion utility available at www.proteomecommons.org (Falkner, Falkner et al. 2007).

### Mass Spectrometry Database Analysis

Merged files were submitted to MASCOT MSMS ion search set to search the published *F. hepatica* genomes (Cwiklinski, Dalton et al. 2015; McNulty, Tort et al. 2017) using the webpage interface (MatrixScience). The following parameters were selected for each peptide search; enzyme set at trypsin with one missed cleavage allowed, fixed modifications set for carbamidomethylation with variable modifications considered for oxidation of methionine, monoisotopic masses with unrestricted protein masses were considered, peptide and fragment mass tolerance were set at ± 1.2 Da and 0.6 Da respectively for an ESI QUAD-TDF instrument (Moxon, LaCourse et al. 2010).

### *In silico* investigation of *Fasciola* transcripts and *F. hepatica* genome

Sequences representing known GST classes were obtained from NCBI (http://www.ncbi.nlm.nih.gov/). A mammalian and a helminth GST sequence were selected for each GST class where available. GST sequences were used to tBLASTn the *F. hepatica* transcriptome (Young, Hall et al. 2010) and the *F. gigantica* transcriptome (Young, Jex et al. 2011) both available to search at (http://bioinfosecond.vet.unimelb.edu.au/wblast2.html). A second *F. hepatica* transcriptome database (EBI-ENA archive ERP000012: an initial characterization of the *F. hepatica* transcriptome using 454-FLX sequencing) was also used to search against. *In silico* investigation of the known GST sequences and positive transcript hit were blasted against genome sequencing project of *F. hepatica* (Cwiklinski, Dalton et al. 2015). Transcript expression levels for individual GST isoforms were analysed from Cwiklinski *et al*. (2018). Each specific GST isoform was used to BLASTp the transcriptome to identify the respective expression level.

### Cloning of newly identified genes

PCR amplification was carried out on an Applied Biosystems 96 Well Thermal Cycler. PCR of cDNA was performed using MyFi Taq (Bioline) following the manufacturer’s instructions. Standard thermocycler conditions involved an initial denaturation at 95°C for two minutes, followed by 25-35 cycles of denaturation (95°C, 30 seconds), annealing (Gradient temperature specific for each gene of interest, 30 seconds) and extension (72°C, 30-90 seconds), before a final extension at 72°C for five minutes and holding period at 4°C until products removed. Primers were based on the scaffolds from the *F. hepatica* genome (FhGST-S2 For: GGGCGATACTATCTATCAACGT Rev: GTGCGACTGACTTTGAATC; FhGST-O2 For: CACACAGCTGGAATTGA TTA Rev: TAATATTGACGGATCCAAACA). PCR products were ligated into pGEM-T-Easy, according to the manufacturer’s protocol and sequenced in house. Sequences were translated using Expasy Translate (https://web.expasy.org/translate/) and molecular weight and pI calculated using Expasy Compute pI/Mw (https://web.expasy.org/compute_pi/). GST domains were predicted using PFam (El-Gebali, Mistry et al. 2019).

### Protein sequence alignment and phylogenetic tree construction

All sequences were aligned using ClustalW through BioEdit Version 7.0.5.3 (10/28/05) (Hall 1999). To construct a phylogenetic tree an alignment of all GST sequences was exported into Molecular Evolutionary Genetics Analysis (MEGA) software version 4.0 (Tamura, Dudley et al. 2007). Analysis was performed using a neighbour-joining method, 1000-replicate, bootstrapped tree. The amino acid data were corrected for a gamma distribution (level set at 1.0) and with a Poisson correction.

### Recombinant *Fasciola hepatica* glutathione transferase Mu class (rFhGST-Mu5) production

FhGST-Mu5 was amplified via PCR using the following primer pair: rFhGST-Mu5 forward primer, **5’ <u>CATATG</u>**GCTCCAGTCTTA 3’; rFhGSTMu5 reverse primer, 5’ **<u>GCGGCCGC</u>**TTAACTGGGTGGTGCA 3’ and a second reverse primer containing the stop codon 5’ **<u>GCGGCCGC</u>**ACTTTAACTGGGTGGTGCA 3’. Restriction enzyme sites (in bold type and underlined) for NdeI (forward primer) and NotI (reverse primer) were included so that the entire ORF could be directly cloned into the pET23a (Novagen) vector. Recombinant proteins were produced in *Escherichia coli* BL21 (DE3) cells (Bioline) as described previously (LaCourse, Perally et al. 2012; Morphew, Eccleston et al. 2012).

*E.coli* preparations containing rFhGST-Mu5 were suspended in lysis buffer (containing 5 mM MgCl, 400 mM NaCl and 20 mM sodium phosphate pH7.4) and were lysed through a freeze/thaw method, freezing in liquid nitrogen followed by thawing at 42°C three times. This was followed by 3 cycles of ultrasonication; the samples were sonicated for 30 seconds with 30 second intervals in ice. The samples were centrifuged at 13,200 x *g* for 20 minutes at 4°C. and purified by GSH-affinity chromatography as described previously.

## Results

### Limited induction of soluble *F. hepatica* GST by TCBZ-SO

Prior to affinity chromatography GST enzymatic specific activity was assessed to examine if overall cytosolic GST activity was induced by TCBZ-SO exposure (Table 1). In general, there was a trend to increased GST specific activity following exposure to TCBZ-SO for treatment groups compared to controls (Supplementary Figure 1). However, following ANOVA no significant difference was noted between any of the treatment groups or the controls (2_df_, F = 1.25, P = 0.320).

**Table 1:** GST specific activity assays of *F. hepatica* GST samples exposed to TCBZ-SO at Control (0 µg/ml), Sub-lethal (15 µg/ml) or Lethal (50 µg/ml) dose. Total activity (nmol/min), total protein (mg) and specific activity (nmol/min/mg) are included.

| Treatment | Total activity (nmol/min). | Total protein (mg). | Specific activity (nmol/min/mg). |
| --- | --- | --- | --- |
| Control | 4463.44 | 8.55 | 522.04 ± 77.92 |
| Sub-lethal | 6002.92 | 10.52 | 570.62 ± 190.46 |
| Lethal | 6190.82 | 9.35 | 662.12 ± 134.63 |

### GST proteomic profiling of individual fluke

Two affinity matrices were then used to isolate GST isoforms from individual adult *F. hepatica*. 18 individuals were homogenised independently and all 18 independently processed through GSH or S-Hexyl GSH agarose columns to separate *F. hepatica* GST proteins from other soluble proteins. Following purification, it was possible to compare the GST-ome from each individual *F. hepatica* exposed to TCBZ-SO, either a Sub-lethal concentration (15 μg/ml) or a Lethal concentration (50 μg/ml), versus those not exposed using 2-DE proteomics.

Proteomic arrays of the GSTs purified from S-hexyl GSH-agarose consistently yielded 13 protein spots (Figure 1A), whereas those purified from GSH-agarose column yielded 11 prominent protein spots (Figure 1B). All protein spots from both purification systems were confirmed as containing *Fasciola* GSTs using tandem mass spectrometry (Table 2; Full proteomic analysis Supplementary Table 1). Comparison of 2-DE protein arrays was then performed to establish if there was a change in abundance of the identified GSTs relating to the different treatment of TCBZ-SO (Figure 1C-F).

**Table 2.**
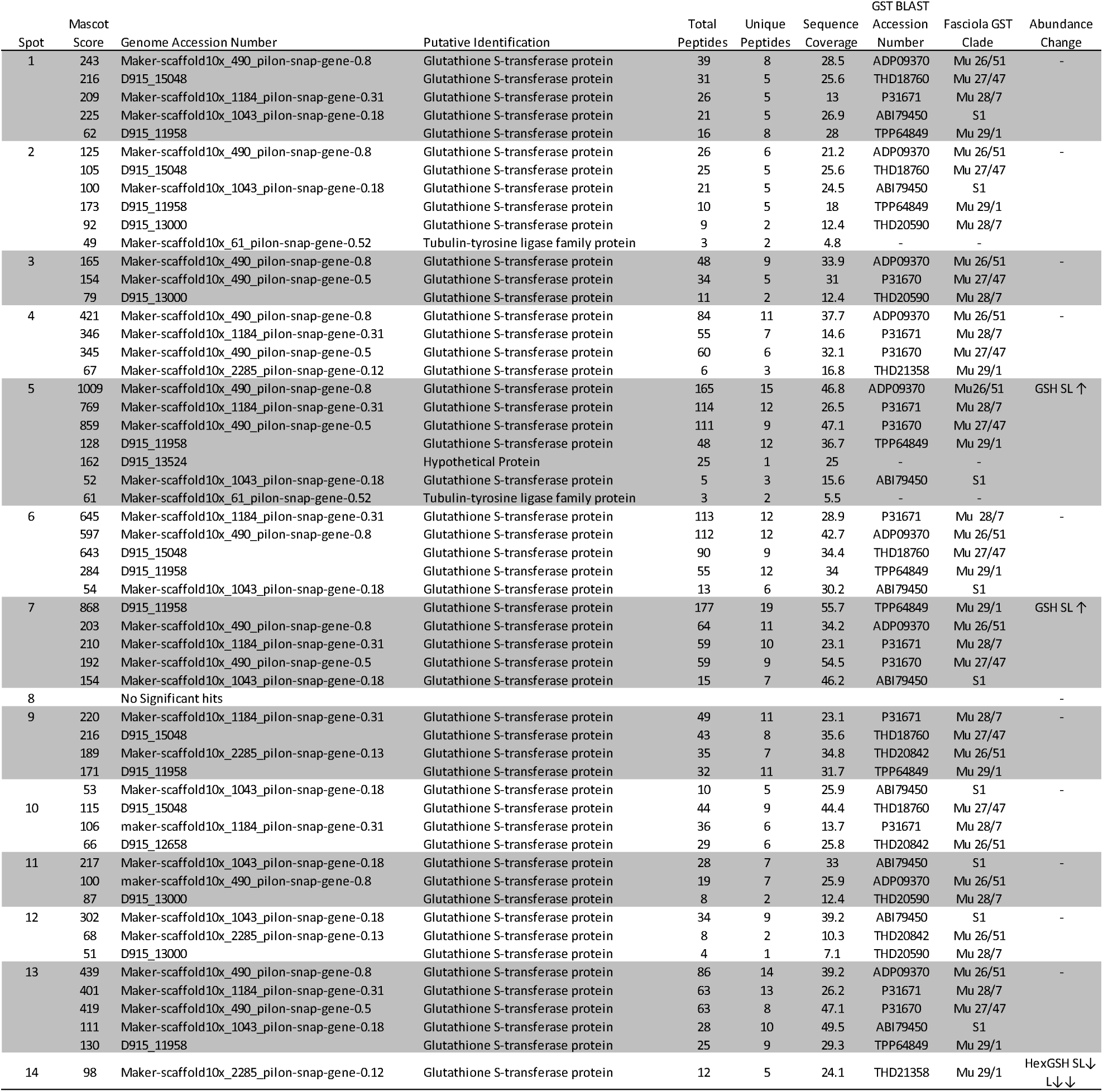
Putative protein identification of GST isoforms from *F. hepatica* by MSMS. Peptide sequences from spots trypsin digested were used to search against both *F. hepatica* genomes for the identification of the specific members of the GST superfamily. MASCOT ion scores of >42 indicate identity or extensive homology (p<0.05). An accession number from Genbank relating to the top scoring BLAST hit to determine GST isoform is also reported. Changes in abundance (↑ or ↓) are denoted for spots responding to sub-lethal or lethal (SL or L) TCBZ-SO exposure for either purification method (GSH or Hex-GSH).

**Figure 1:**
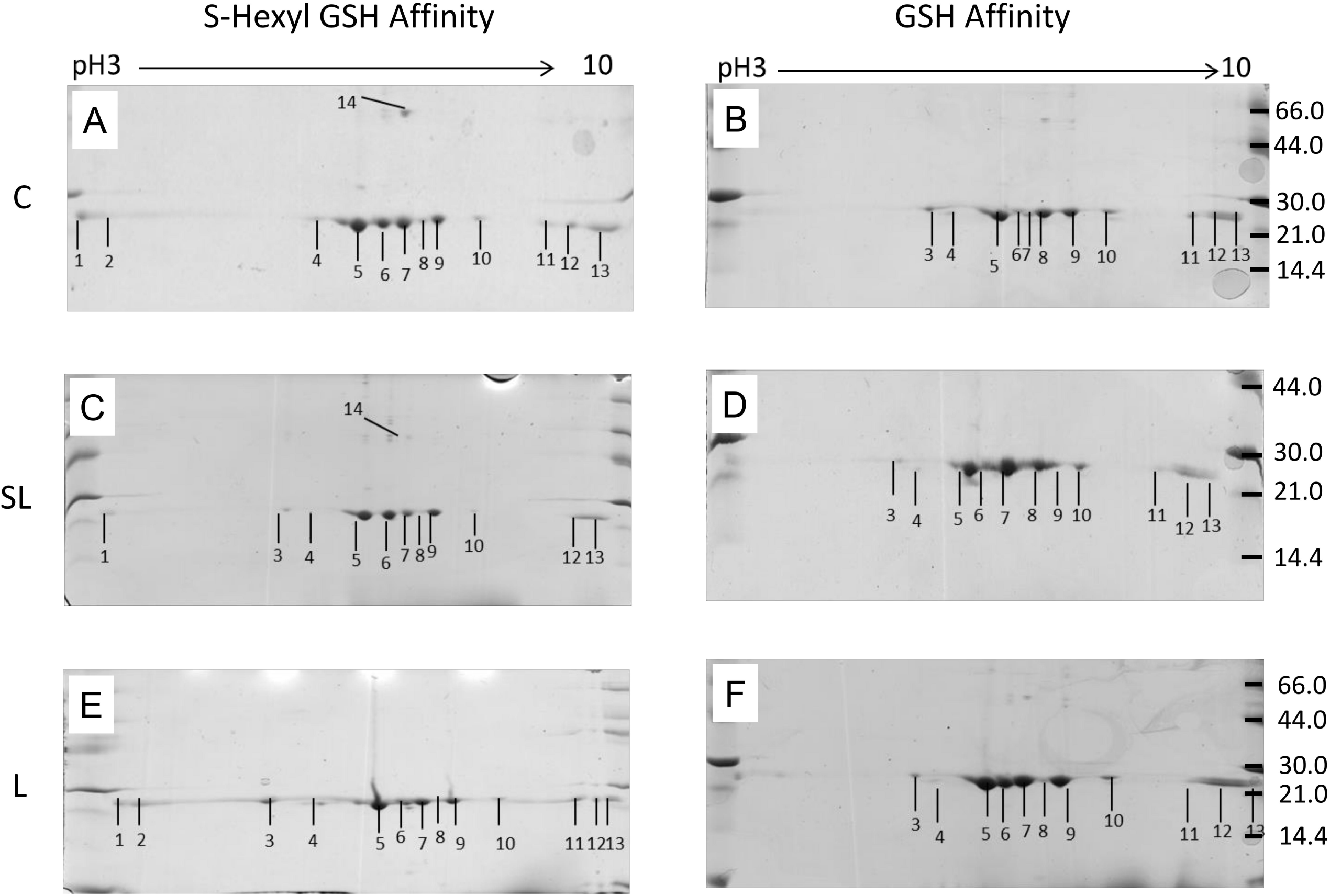
Representative 2-DE arrays of GSTs purified from *F. hepatica* using S-hexyl GSH and GSH agarose columns following TCBZ-SO exposure. A) S-hexyl GSH agarose purified GSTs from Control samples (TCBZ-SO 0 µg/ml). B) GSH agarose purified GSTs from Control samples (TCBZ-SO 0 µg/ml). C) S-hexyl GSH agarose purified GSTs from Sub-lethal samples (TCBZ-SO 15 µg/ml). D) GSH agarose purified GSTs from Sub-lethal samples (TCBZ-SO 15 µg/ml). E) S-hexyl GSH agarose purified GSTs from Lethal samples (TCBZ-SO 50 µg/ml). F) GSH agarose purified GSTs from Lethal samples (TCBZ-SO 50 µg/ml). All arrays were run on 7 cm IPG strips pH 3-10, 12.5% SDS PAGE and Coomassie blue stained. Spot numbers relate to GST putative identifications seen in Table 2.

When comparing the S-Hexyl GSH Control array with both the S-Hexyl GSH TCBZ-SO exposed arrays it was noted that spot 14 (Figure 1A,C and E) was present on all Control arrays, thus present on the average Control array. However, the presence of this protein spot varied on the TCBZ-SO treatment arrays (Sub-lethal & Lethal). This particular protein spot was only present on 2 Sub-lethal arrays and 1 Lethal array. MSMS analysis identified this spot as Mu class GST29 (THD21358).

The comparison of the arrays produced via GSH agarose affinity columns identified two protein spots of interest when the average Control and average Sub-lethal arrays were examined with both spots increased in abundance in Sub-lethal samples; Spot 5, Mu class GST 26, (1_df_, F = 3.89, P = 0.089) and Spot 7, Mu class GST 29, (1_df_, F = 4.83, P = 0.064) both approaching statistical significance (Figure 1B and D; Supplementary Figure 2). No additional changes in protein abundance were observed for GSH purifications.

### GST expression in the cytosol of individual fluke and affinity binding

Given the potential of Mu class GSTs responding to TCBZ-SO exposure, Western blotting was used to estimate the number of Mu class GSTs present in liver fluke cytosol prior to affinity chromatography, given previous indications that some GST isoforms may fail to bind to affinity matrices (Brophy, Crowley et al. 1990). Assays were undertaken on five individual adult fluke using anti-*S. mansoni* Mu class polyclonal antibodies, previously shown to recognise *F. hepatica* Mu class GSTs (Chemale, Morphew et al. 2006). The anti-flatworm GST Mu class antibody recognised 8 GST subunits within the cytosolic profile (Figure 2A). Post purification the blot patterns display the same distinctive GST protein profiles following both GSH and S-Hexyl GSH affinity 2-DE gels (Figure 2B-C). A distinctive and reproducible 2-DE GST profile provides evidence that 8 GST subunits are recognized by the Mu antibody post purification.

**Figure 2:**
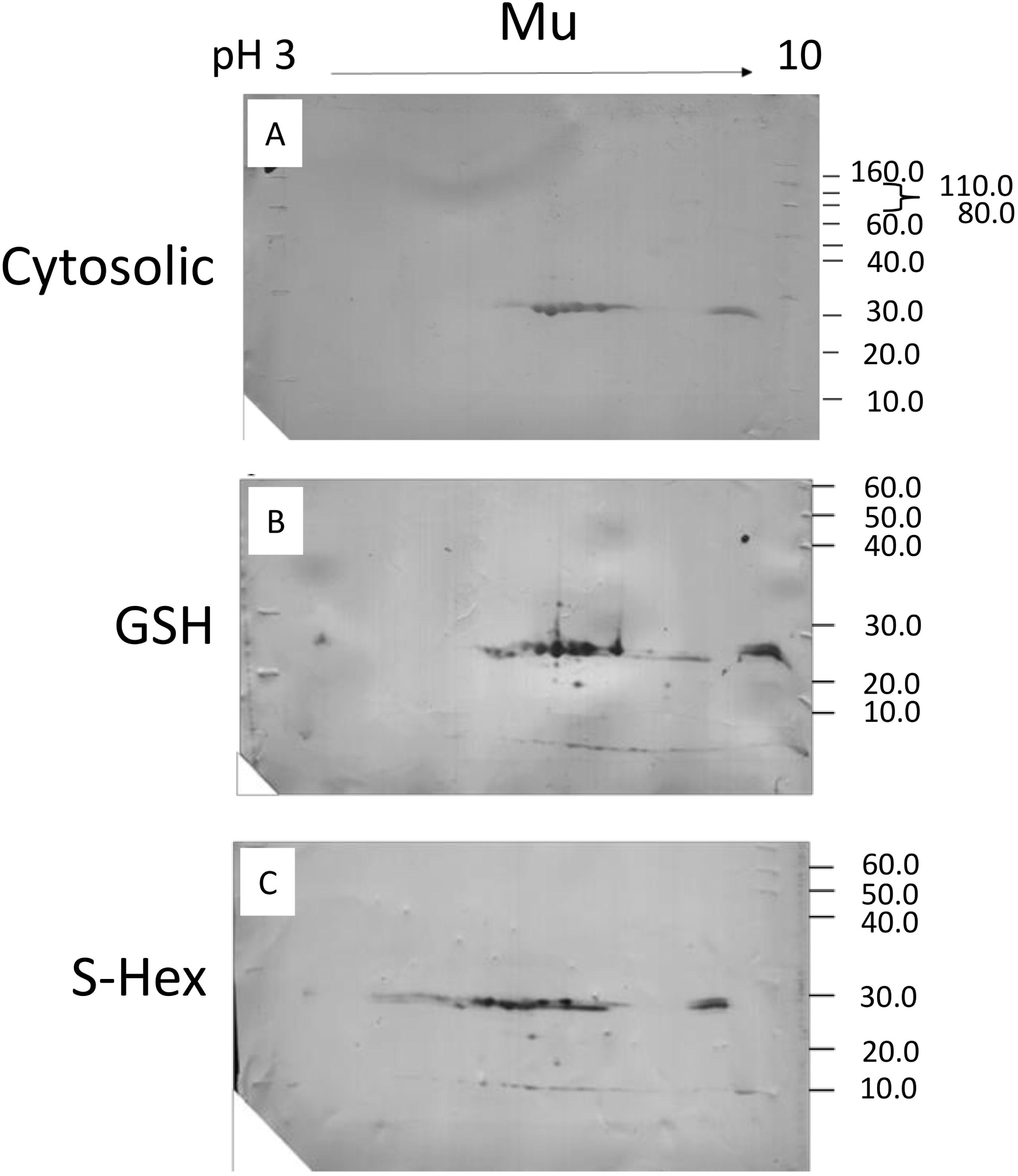
Assessment of Mu class GST binding affinity through Western blotting of affinity purified GSTs comparison to cytosolic fractions. A) Visualisation of TCA precipitated cytosolic proteins of *F. hepatica* adult worms using two-dimensional gel electrophoresis (2-DE) and Western blot analysis probing for Mu class GSTs. In total, 100 µg of cytocolic protein was resolved on non-linear IPG strips and 12.5% polyacrylamide gels. B) and C) Visualisation of GSH agarose purified GST subunits of *F. hepatica* adult worms using two-dimensional gel electrophoresis (2-DE) and Western blot analysis analysis probing for Mu class GSTs. In total, 5 µg of GSH or S-Hexyl GSH purified GSTs were resolved on non-linear IPG strips and 12.5% polyacrylamide gels. A), B) and C) Western blots probed with anti-*S. mansoni* Mu antibody (1:5,000 dilution) and developed using alkaline phosphatase linked secondary antibody (anti-Goat IgG).

### Bioinformatic characterisation of GSTs identified in *F. hepatica*

Following analysis of available transcript and genome sequences the known 4 Mu class GSTs were identified alongside a fifth Mu class GST designated FhGST-Mu5. Following cloning and sequencing of FhGST-Mu5, multiple alignment of all Mu class GSTs of *F. hepatica* revealed the extent of identity and similarity across this class of GSTs (Figure 3A). Amino acid sequence similarity when comparing the newly identified FhGST-Mu5 (Genbank MT613329) with the previously known *F. hepatica* Mu class GSTs identified the closest sequence similarity was with FhGST-7 at approximately 54%. It is also worth noting that Genbank entry THD26413 matches to FhGST-Mu5 with 91.9% sequence identity but is an incomplete sequence lacking the N-terminus. When transcript expression was analysed for FhGST-Mu5 based on Cwiklinski *et al*. (2018), the levels of transcript within the adult is significantly lower than in alternative life cycle stages such as metacercariae and newly excysted juveniles from 1 to 24 h (Figure 3B).

**Figure 3.**
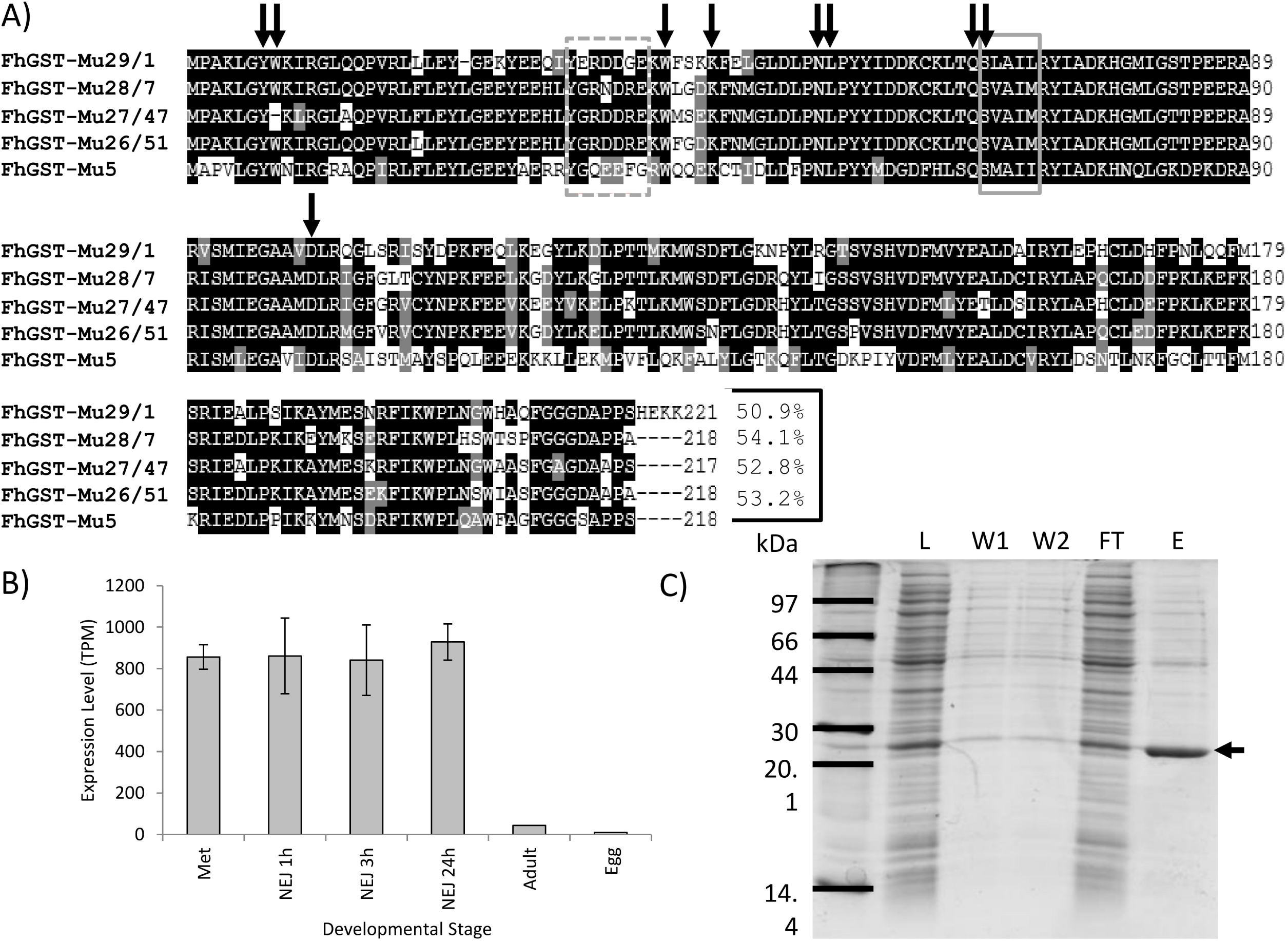
Bioinformatics, Expression and purification of recombinant rFhGST-Mu5. A) Multiple alignment of the 4 established *F. hepatica* Mu class GSTs and the newly identified FhGST-Mu5 revealed through transcriptome/genome analysis. No other Mu class GSTs were identified within the genome of *F. hepatica*. The consensus sequence SNAIL/TRAIL and their synonymous sequences in parasites are in the solid-line grey box. The residues forming the µ-loop are in dotted-line grey box. Arrowed are predicted GSH binding sites. Amino acid sequence identity of FhGST-Mu5 with the four previously known Mu class GSTs is provided at the end of the alignment. Accession numbers for each Mu class GST used: FhGST-Mu29/1 (P56598), FhGST-Mu28/7 (P31671), FhGST-Mu27/47 (P31670) and FhGST-Mu26/51 (P30112). B) Transcript expression levels for FhGST-Mu5 were analysed from Cwiklinski *et al*. (2018). C) SDS-PAGE gel of the expression and purification of rFhGST-Mu. L: *E. coli* total cytosolic protein lysate, 10 µg. W1 and W2: Column washes removing non-binding proteins, 10 µl. FT: Flow through proteins collected after passing through a GSH-agarose column, 10 µg. E: Eluted GSH-affinity purified recombinant rFhGST-Mu5 protein, 2 µg. Run on 12.5% SDS-PAGE and Coommassie blue stained. Arrowed is the band representing rFhGST-Mu5.

Further transcript and genome investigation allowed the examination of the complete GSTome of *F. hepatica*. In addition to FhGST-Mu5, *in silico* investigation revealed the identification of a second Sigma class and a second Omega class GST. Bioinformatic characterization of the new FhGST-S2 and FhGST-O2 was undertaken to identify the structural features and characteristics of these genes/proteins. Only a single homologue for each was identified in the original *F. hepatica* genome. For FhGST-S2, gene BN1106_s1104B000225 (Scaffold 1104) was identified yet this is now fragmented and incomplete in the most recent version of the genome (PRJEB25283) despite transcript support (Supplemental Table 2). For FhGST-O2, gene BN1106_s50B000678 (Scaffold 50) was revealed and is now designated as maker-scaffold10x_938_pilon-snap-gene-0.52/D915_03058. Each gene encoded for a predicted single protein isoform.

Both the newly predicted FhGST-S2 and FhGST-O2 were cloned (Supplemental Figure 3A) and sequenced. Confirmation of the correct class assignment was performed with multiple alignment (Supplemental Figure 3B and 3C) and comparison of gene intron exon structure (Supplemental Figure 4). Of note was a significant N-terminal extension of 20 amino acids in FhGST-S2 when compared to FhGST-S1. FhGST-O2 in comparison to FhGST-O1 revealed the addition of 1 amino acid to each of exons 1 and 5. Further confirmation of class assignment was support with both FhGST-S2 and FhGST-O2 subjected to a PFam domain analysis revealing key predicted GST features; FhGST-S2 with a predicted C-terminal domain (PFam GST_C_3) and FhGST-O2 with a predicted N- and C-terminal domain (PFam GST_N_3 and GST_C_2).

Following a full phylogenetic analysis of the completed *F. hepatica* GST-ome, all of the newly identified FhGST-Mu5, FhGST-S2 and FhGST-O2 were assigned to their respective clades (Figure 4). Of note is the close association of FhGST-Mu5 to the Schistosome Mu class GSTs rather than to the previously established four *Fasciola* Mu class isoforms.

**Figure 4:**
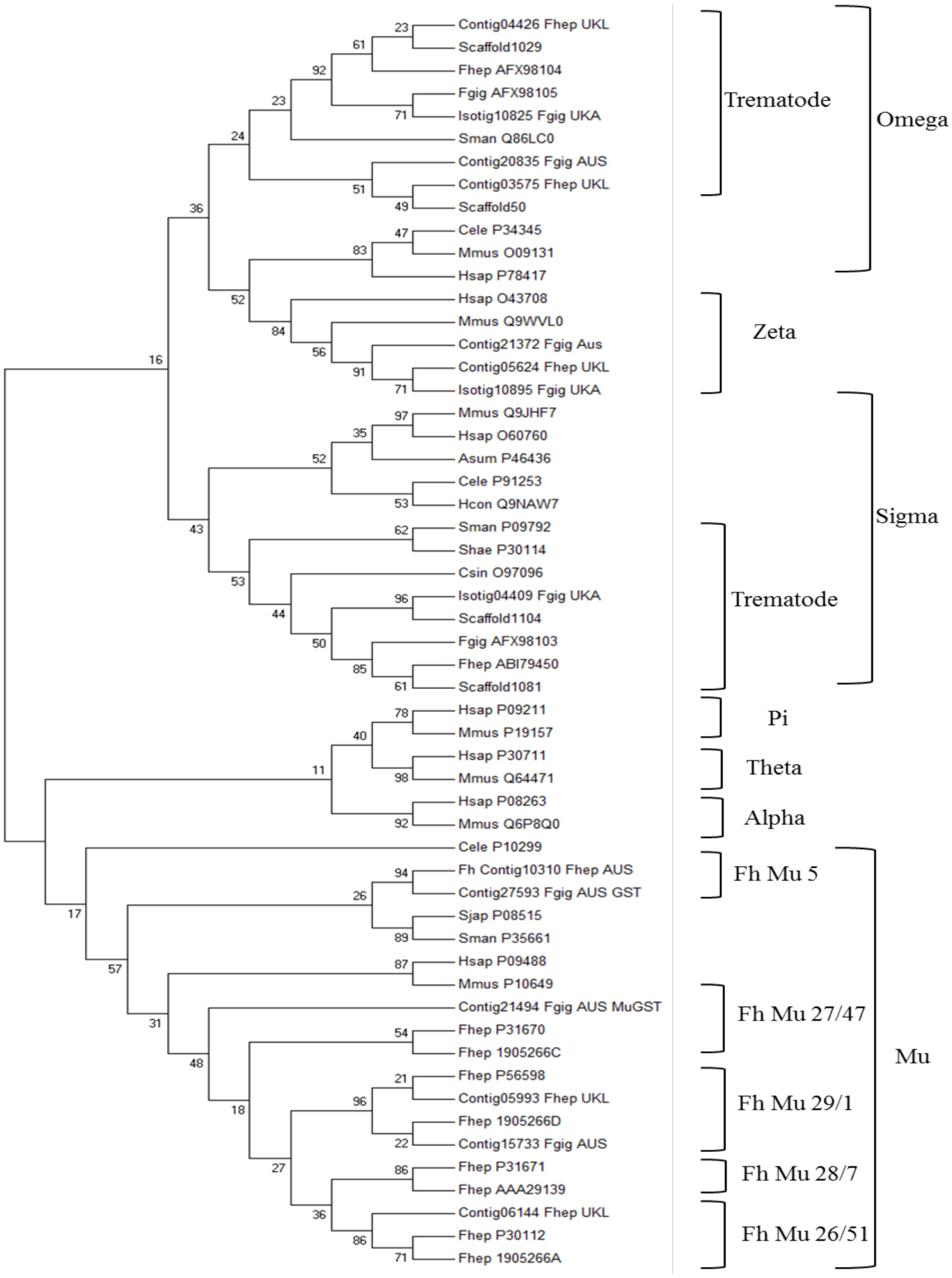
Phylogenetic analysis of the soluble cytosolic GST superfamily. Neighbour-joining phylogenetic tree constructed using amino acid sequences through MEGA v 7.0 with 1000 bootstrapped support and a Poisson correction. All reported accession numbers are from Genbank. Where sequences were identified *in silico*, only contig numbers are reported. Those from *F. gigantica* were taken from the study of Young *et al*. (2011) and transcripts produced by Aberystwyth University. Those from *F. hepatica* were taken from the study of Young *et al*. (2010) and transcripts produced by the University of Liverpool (EBI-ENA archive ERP000012: An initial characterization of the *F. hepatica* transcriptome using 454-FLX sequencing).

### Expression, purification and characterisation of rFhGST-Mu5

Full sequence length recombinant *F. hepatica* Mu class GST (rFhGST-Mu5) was expressed and purified from transformed *E.coli* cytosol following expression in BL21 (DE3) cells. Purity was assessed on SDS-PAGE gels (Figure 3C). Interestingly, rFhGST-Mu5 was not able to be produced as a pure protein with significant levels of contaminating *E. coli* proteins remaining in the sample following GSH affinity purification. However, rFhGST-Mu5 was produced as an active protein for further studies displaying enzymatic activity towards the model GST substrate 1-chloro-2,4-dinitrobenzene (CDNB). The specific activity for the rFhGST-Mu5 preparation was confirmed at 243.27 ± 92.45 nmol/min/mg.

## Discussion

The 2-DE mapping of GSTs has been shown to be a useful tool to delineate the function of individual members of this soluble protein superfamily (Chemale, Morphew et al. 2006; Morphew, Eccleston et al. 2012), particularly as these proteins play a role in Phase II detoxification (Cvilink, Lamka et al. 2009). To date, research has only been completed on pooled cytosol samples from wild-type fluke and defined isolates and there has not been a robust sub-proteomic study that compared the expression of GST isotypes in individual fluke populations under TCBZ-SO challenge in culture. This study has adapted the pooled approach and, for the first time, performed analytical scale 2-DE mapping of GSTs from individual *F. hepatica* adult parasites. Thus, GSTs were purified from the cytosol of single adult flukes using either S-Hexyl Glutathione or Glutathione agarose columns, resolved using analytical 2-DE and identified individual GSTs by MSMS with the support of liver fluke transcriptomic and genomic databases. In doing so, we can identify individual fluke responses within the GST superfamily following exposure to chemotherapeutics. Furthermore, this finding has major implications for future population and resistance monitoring studies specifically on, but not limited to, liver fluke GSTs.

In the current study, both S-Hexyl GSH agarose and GSH agarose columns were used for GST purification at the individual fluke level. Previous studies (Chemale, Morphew et al. 2006; Morphew, Eccleston et al. 2012) have demonstrated that S-Hexyl GSH agarose columns have the ability to purify a greater range of GSTs in both *F. hepatica* and *F. gigantica* population mixes respectively thus was a useful inclusion in the current work at the individual fluke level. Using biochemical techniques and analytical sub-proteomics identified both Sigma and Mu class GSTs purified from individual adult *F. hepatica*. It was confirmed that both S-Hexyl GSH and GSH agarose columns have the ability to purify both Mu and Sigma class GSTs, but with GSH columns purifying the Sigma class to a much lesser extent expressing a preference to purify Mu class GSTs as observed for pooled samples (Chemale, Morphew et al. 2006).

The overall GST-ome profile, via GST activity and 2-DE arrays, demonstrated a general trend of response to TCBZ-SO exposure. Following exposure GST specific activity increased with increasing TCBZ-SO concentration. In addition, the only changes noted in both S-Hexyl GSH and GSH agarose purifications were recorded abundance changes associated with Mu class GSTs, specifically FhGST-Mu29 and FhGST-Mu26. Therefore, from activity data and proteomic profiling, it is likely that, of the two GST classes identified, Mu class GSTs are likely highly important for xenobiotic detoxification with Sigma class GSTs acting as secondary xenobiotic sequesters with a primary role as a house-keeping enzyme and as, more importantly, an immunomodulatory (LaCourse, Perally et al. 2012). This finding of Mu class GST-TCBZ-SO detoxification supports the work of Chemale *et al*. (2010) examining the TCBZ-SO response of TCBZ-resistant and TCBZ-susceptible isolates. In this case, both FhGST-Mu29 and FhGST-Mu26 responded to TCBZ-SO exposure, in agreement with the current study. We identified changes in response to TCBZ-SO exposure linked to dimer and monomer formation of FhGST-Mu29 and differential purification of both using the two purification methods. S-hexyl GSH purification was more efficient at purifying FhGST-Mu29 dimers compared to GSH agarose purification. On exposure to TCBZ-SO a reduction in FhGST-Mu29 dimers was observed with a corresponding increase in FhGST-Mu29 monomers purified through GSH agarose. The novel dimer-monomer GST conformational switch might reflect a new liver fluke mechanism in response to TCBZ-SO challenge. GSTs normally function as dimers but active monomeric GSTs have been previously identified in *F. hepatica* (Brophy et al., 1990).

In *F. hepatica* there are four recognised isoforms of Mu class GSTs i.e. FhGST-Mu26, 27, 28 & 29 (alternatively called FhGST-Mu51, 47, 7 & 1) with a fifth identified only through bioinformatics previously (Morphew, Eccleston et al. 2012), and now cloned and expressed in the current work. Alongside the identification of FhGST-S1, four of the five Mu class GST isoforms were identified in the samples examined in the current study under TCBZ-SO stress. In previous proteomic studies the same four classes have also been identified. However, the functional significance of multiple Mu GSTs is as yet unknown. Multiple Mu class isoforms might relate to their role in the protection of the parasite from various classes of xenobiotics derived from the host bile environment (Brophy, Mackintosh et al. 2012). Specifically, the current work supports a role for FhGST-Mu29 in TCBZ-SO response via conformational changes as identified by evidence of altered in dimer/monomer ratios. Of interest, based on transcriptome evidence of Cwiklinski *et al*. (2018) FhGST-Mu29 is naturally the highest expressed Mu class GST in adult fluke. In addition, FhGST-Mu26 ranks third in all Mu class GST expression (FhGST-Mu29 > Mu27 > Mu26 > Mu5 > Mu28). Therefore, it is likely that these primary expressed GSTs are important in binding xenobiotics with structures such as such as TCBZ-SO.

In many cases peptides belonging to different GSTs were identified in a single protein spot providing the identification of multiple GST isoforms. As reported by Chemale *et al*. (2006), this may result from spot overlapping in the 2-DE gels, as proteins may have a similar p*I*, potential modifications and co-migration. Of note is the failure to identify the fifth Mu class GST, FhGST-Mu5, despite overlapping GST isoforms identified in multiple spots. Given a sequence similarity of 54% for FhGST-Mu5 compared to FhGST-7 the failure to identify FhGST-Mu5 is unlikely to be from miss assigning sequenced peptides to alternative Mu class GSTs and likely represents low expression as evidenced from transcriptomics (Cwiklinski, Jewhurst et al. 2018) or, given the poor affinity purification of FhGST-Mu5, non-binding to affinity columns.

In an attempt to assess if FhGST-Mu5 was not identified in affinity purified samples as a result of non-binding, *F. hepatica* cytosolic material was probed with anti-*S. mansoni* Mu polyclonal antibodies and compared with the profiles obtained post affinity purification. Given that the same repertoire of protein spots following Western blotting was visualised on both cytosolic and affinity purified fractions, in addition to FhGST-Mu5 recognition by anti-*S. mansoni* Mu (data not shown), it seems unlikely that FhGST-Mu5 was missed in the affinity proteomics study. In addition, it seems unlikely that FhGST-Mu5 was missed due to low expression in adults given the identification of FhGST-Mu28 in the current work and in previous studies (Chemale, Morphew et al. 2006). Thus, the potential exists that FhGST-Mu5 is a low affinity isoform. In support, Brophy *et al*. (1990) proposed that an endogenous ligand interacts with GSTs preventing GSTs binding to the affinity matrix generating a ‘low affinity’ fraction. Therefore, general inhibitory binding factors are likely present in the liver fluke cytosol and may be important in flatworm GST function.

Following the successful induction and expression of FhGST-Mu5 it is clear that ‘low affinity’ GSTs are produced within the GST-ome of *F. hepatica* yet not all GSTs fail to bind from potential inhibitory factors. GSH affinity purification of rFhGST-Mu5 resulted in low impure yields of recombinant protein and suggests that FhGST-Mu5 is a ‘low affinity’ isoform. Previous studies have all successfully used glutathione affinity chromatography for successful purification of native and recombinant GSTs (Chemale, Morphew et al. 2006; LaCourse, Perally et al. 2012; Morphew, Eccleston et al. 2012) yet failed to purify FhGST-Mu5. Thus, to determine if rFhGST-Mu5 is an isoform with ‘low affinity’ for glutathione the specific activity was determined with the model substrate CDNB (Habig, Pabst et al. 1974). The specific activity of rFhGST-Mu5 was significantly lower than that recorded for the previously known 4 Mu Class GSTs from *F. hepatica* (Salvatore, Wijffels et al. 1995; Kalita, Shukla et al. 2017). This lower affinity may be correlated with the lower sequence homology and the more distant grouping of FhGST-Mu5 in phylogenetic modelling aligning closer to schistosome Mu class GSTs rather than the previous four *F. hepatica* Mu class. Brophy *et al*. (1990) demonstrated that following chromatofocusing 95% of ‘low affinity’ GSTs were relieved of their inhibition and thus, based on current evidence, it is likely that FhGST-Mu5 could indeed be classed as a ‘low affinity’ Mu class GST as part of the remaining 5% of activity. Low GSH affinity most likely accounts for the previous lack of detection during affinity studies with the initial identification achieved through transcriptomic analysis (Morphew, Eccleston et al. 2012). Given that FhGST-Mu5 clustered with schistosome Mu class GSTs during phylogenetics it is possible that FhGST-Mu5 and schistosome Mu class GSTs perform similar roles within these fluke species.

The current study represents the first 2-DE profiling of TCBZ-SO exposed *F. hepatica* GSTs. However, TCBZ-SO stress in *F. gigantica*, and the resulting GST activity, has been previously investigated. Shehab *et al*. (2009) examined GST activities from crude homogenates of adult and juvenile *F. gigantica* exposed to TCBZ-SO concentrations. This research indicated that a significant increase in the level of GST was present, in both adult and juvenile flukes, after exposure to TCBZ-SO (Shehab, Ebeid et al. 2009). Such a significant increase in response to TCBZ-SO prior to affinity purification was not noted in the current research and may reflect important differences between *F. hepatica* and *F. gigantica* GST expression. Nevertheless, the work of Shehab and colleagues further supports the role of Mu class GST in TCBZ-SO detoxification.

The release of the genome assemblies of *F. hepatica* (Cwiklinski, Dalton et al. 2015; McNulty, Tort et al. 2017) has allowed for further in-depth and complete investigation of the GST-ome complement of this parasitic flatworm. Two new soluble superfamily GSTs were identified; a second Sigma (σ) class and a second Omega (ω) class, on original genes BN1106_s1104B000225 and BN1106_s50B000678 (scaffolds 1104 and 50, respectively). Both GSTs contained Pfam IDs for the respective GSTs and both sequences where successfully amplified through PCR and sequence verified. The predicted molecular weight of the sub-units of the newly identified Sigma and Omega GSTs were shown to be 26 and 27 kDa respectively, and this is in general agreement with known soluble GSTs that have a subunit mass of between 23 and 28 kDa with an average length of 220 amino acids (Torres-Rivera and Landa 2008). Gene structure analysing introns and exons for both the newly identified Sigma and Omega genes in comparison with the previously identified *F. hepatica* Sigma and Omega supported the confirmation of GST class assignment.

Previous research has demonstrated that model organisms (humans and mice) both encode for 2 Omega class GST genes which are widely expressed (Board 2011) reflecting expression within *F. hepatica*, albeit human and mice omega GSTs comprise of six exons (Board 2011) rather than 5 in *F. hepatica* omega class GSTs. Interestingly, omega class GSTs have been linked with drug resistance in human cancers (Townsend and Tew 2003) and Alzheimer’s disease (Allen, Zou et al. 2012) and thus may have some role in anthelmintic resistance or detoxification not yet discovered.

Sigma class GSTs in *F. hepatica* were also initially identified by Chemale *et al*. (2006). A recombinant form of *F. hepatica* Sigma class GST, FhGST-S1, has since been produced and demonstrated to have multi-functional roles, including general endogenous detoxification, and is strongly linked with prostaglandin synthesis and the modulation of dendritic cell activity (LaCourse, Perally et al. 2012). Across trematode species the exon-intron structure of Sigma class GSTs is conserved. Recently, reports of 5 newly identified Sigma class GSTs from *Clonorchis sinensis* consist of 4 exons akin to the two *F. hepatica* genes (Bae, Kim et al. 2016). It was also noted that the final exon, exon 4, of Sigma GST genes in the gene predictions of all the trematode species investigated by Bae *et al*. (2016) consisted of 225 bp; this conservation of gene structure likely reflects conserved biological function. As yet, proteomic investigations have not identified FhGST-S2 from adult flukes despite their presence in adult transcriptomes. It is therefore likely that FhGST-S2 remains part of the unbound fraction of the GST-ome; a likely ‘low affinity’ sigma class GST.

With a key role for GSTs in the detoxification of TCBZ demonstrated through proteomic profiling it is now crucial to understand any involvement of GSTs in TCBZ resistance. This is of particular importance given that Scarcella *et al*. (2012) identified that fluke resistant to TCBZ expressed significantly higher levels of GST activity compared to susceptible flukes. The authors suggest that under TCBZ-SO exposure there is an increased requirement for Phase I detoxification of TCBZ-SO, to the less effective TCBZ-SO_2_, and thus also require increased Phase II detoxification, principally from GSTs, to catalyse TCBZ intermediates. Given the recent bioinformatics identification of a potential Cytochrome P450 (Cwiklinski, Dalton et al. 2015), TCBZ-SO exposure is likely to stimulate this Phase I pathway leading to an increased requirement for phase II GSTs. Therefore, profiling the specific GST isoforms will give more insight into resistance mechanisms.

## Conclusions

Glutathione transferases (GSTs) are a multi-gene family of ubiquitous multifunctional proteins that are predicted to have major roles in detoxifying both endogenous and exogenous toxins as part of the Phase II system. We have expanded the knowledge on this important protein family in the parasitic flatworm *F. hepatica*. In doing so, we have revealed novel GST members including ‘low affinity’ Mu and Sigma class enzymes. In addition, it is clear that GSTs respond to TCBZ-SO exposure and the role of GSTs in TCBZ resistance awaits further investigation. Finally, the ability to incorporate individual fluke for proteomic and sub-proteomic studies has implications for potential early TCBZ resistance monitoring in liver fluke populations.

## Acknowledgements

The authors are grateful to Randall Parker Foods (Wales) for providing *F. hepatica* infected sheep livers. This work was supported by the Biotechnology and Biological Sciences Research Council through an IBERS PhD Scholarship award and through Innovate UK (Grant Number: 102108).

