## Supplemental Figure and Table Legends and Supplemental Table 2 for "The soluble glutathione transferase superfamily: Role of Mu class in Triclabendazole sulphoxide challenge in *Fasciola hepatica*"

**Supplemental Figure 1**: Glutathione transferase specific activity (nmol/mg/min) assessed with the model substrate CDNB for *F. hepatica* cytosolic samples for each TCBZ-SO treatment. Treatment groups, control, sub-lethal, and lethal, contained adult fluke that were exposed to TCBZ-SO at 0, 15 and 50 µg/ml respectively.

**Supplemental Figure 2**: Comparison of the 2-DE arrays for each TCBZ-SO treatment group via Progenesis. Comparison of the average Control (A) vs. average Sub-lethal treatment (B) purified via GSH agarose identified an increase in normalised spot volumes for two spots, spots 5 and 7. Comparison of the average Control (C) vs. average Lethal (D) purified via GSH agarose identified an increase in normalised spot volumes for spot 5. Proteins were separated across a linear pH range 3 -10 using IEF in the first dimension and 12.5% SDS-PAGE in the second dimension and Coomassie Blue-stained

**Supplemental Figure 3**: Bioinformatics and cloning of the newly identified FhGST-S2 and FhGST-O2. A) PCR amplification of FhGST-O2 and FhGST-S2: 1% agarose gel displaying full PCR products of FhGST-O2 (Lane 2 and 3) and FhGST-S2 (lane 4 & 5). Negative control in lane 6. Replicate lanes represent different individual cDNAs. Both products, for FhGST-O2 and FhGST-S2, were subsequently cloned into pGEM-T-Easy and sequenced in house. B) Multiple alignment of *Fasciola* Sigma class GST protein sequences with those of genome sequenced organisms. All GST sequences were aligned using Clustal W. The amino acids involved in the formation of glutathione-binding and catalytic sites are marked with a plus (+) and asterisks (*), respectively. C) Multiple alignment of *Fasciola* Omega class GST protein sequences with those of genome sequenced organisms. All GST sequences were aligned using Clustal W. Indicated by a solid line are the proline-rich residues in the Omega class characteristic N-terminal extension. The catalytic cysteine residue characteristic of Omega class GSTs is indicated with *. Residues indicated with + relate to the identified Omega class GST motifs identified by Chemale *et al*. (2006). Species abbreviations used are Mmus, *Mus musculus*; Hsap, *Homo sapiens*; Asum, *Ascaris suum*; Cele, *Caenorhabditis elegans*; Hcon, *Haemonchus contortus*; Sman, *Schistosoma mansoni*; Shae, *Schistosoma haematobium*; Csin, *Clonorchis sinensis*.

**Supplemental Figure 4:** Comparison of exon-intron architectures of sigma and omega class GST genes from *F. hepatica*. Protein-coding regions are shown as boxes, and intervening introns are demonstrated by solid thin lines. The lengths of exons and introns in base pairs are presented at the corresponding positions. Of note is the N terminal extension in exon 1 of FhGST-S2. FhGST-S1 (Scaffold 1081), FhGST-S2 (Scaffold 1104), FhGST-O1 (Scaffold 1029) and FHGST-O2 (Scaffold 50).

**Supplementary Table 1.** Full details of the putative protein identification of GST isoforms from *F. hepatica* by MSMS. Peptide sequences from spots trypsin digested were used to search against both *F. hepatica* genomes for the identification of the specific members of the GST superfamily. MASCOT ion scores of >42 indicate identity or extensive homology (p<0.05). An accession number from Genbank relating to the top scoring BLAST hit to determine GST isoform is also reported.

**Supplemental Table 2:** Transcript support for the newly identified FhGST-S2 and FhGST-O1. BLAST hits following analysis of databases using FhGST-S1, -S2, -O1 and -O2 are provided. Top scoring hits are shaded in grey. Transcripts were retrieved from ^A^Young *et al*. (2011), ^B^Young *et al*. (2010), ^C^an in house *F. gigantica* newly excysted juvenile transcriptome, ^D^the available EBI-ENA archive ERP000012: an initial characterization of the *F. hepatica* transcriptome using 454-FLX sequencing and ^E^from *F. hepatica* ESTs available by anonymous FTP from the Wellcome Trust Sanger Institute <ftp://ftp.sanger.ac.uk/pub/pathogens/Fasciola/>.

| **Sigma Class GSTs** | **GST-S1** | **GST-S2** |
| --- | --- | --- |
|  | **Scaffold1081** | **Scaffold1104** |
| Contig24647^A^ | 1.00E-33 | 6.00E-17 |
| Contig27045^A^ | 2.00E-10 | - |
| Fh_Contig8894^B^ | 4.00E-50 | 5.00E-24 |
| isotig05100^C^ | 8.00E-50 | 2.00E-23 |
| isotig05101^C^ | 8.00E-50 | 2.00E-23 |
| isotig04409^C^ | 6.00E-14 | 5.00E-50 |
| contig07874^D^ | 6.00E-50 | 1.00E-23 |
| contig06001^D^ | 7.00E-14 | 6.00E-50 |
| **Omega Class GSTs** | **GST-O1** | **GST-O2** |
|  | **Scaffold1029** | **Scaffold50** |
| contig03575^D^ | 3.00E-11 | 2.00E-45 |
| contig04426^D^ | 2.00E-45 | 2.00E-11 |
| HAN5016c12.q1kT3^E^ | 5.00E-04 | 5.00E-24 |
| isotig09504^C^ | 1.00E-10 | 3.00E-44 |
| isotig10825^C^ | 1.00E-42 | 9.00E-12 |
| Contig20835^A^ | 1.00E-10 | 3.00E-44 |
| Contig13488^A^ | 5.00E-35 | 4.00E-09 |
| Contig12260^A^ | 3.00E-43 | 2.00E-12 |
| Fh_Contig5261^B^ | 2.00E-45 | 2.00E-11 |
| Fh_Contig2859^B^ | 6.00E-11 | 1.00E-44 |
