## Supplementary figures and images for "The soluble glutathione transferase superfamily: Role of Mu class in Triclabendazole sulphoxide challenge in *Fasciola hepatica*"

### Supplemental Figure 1

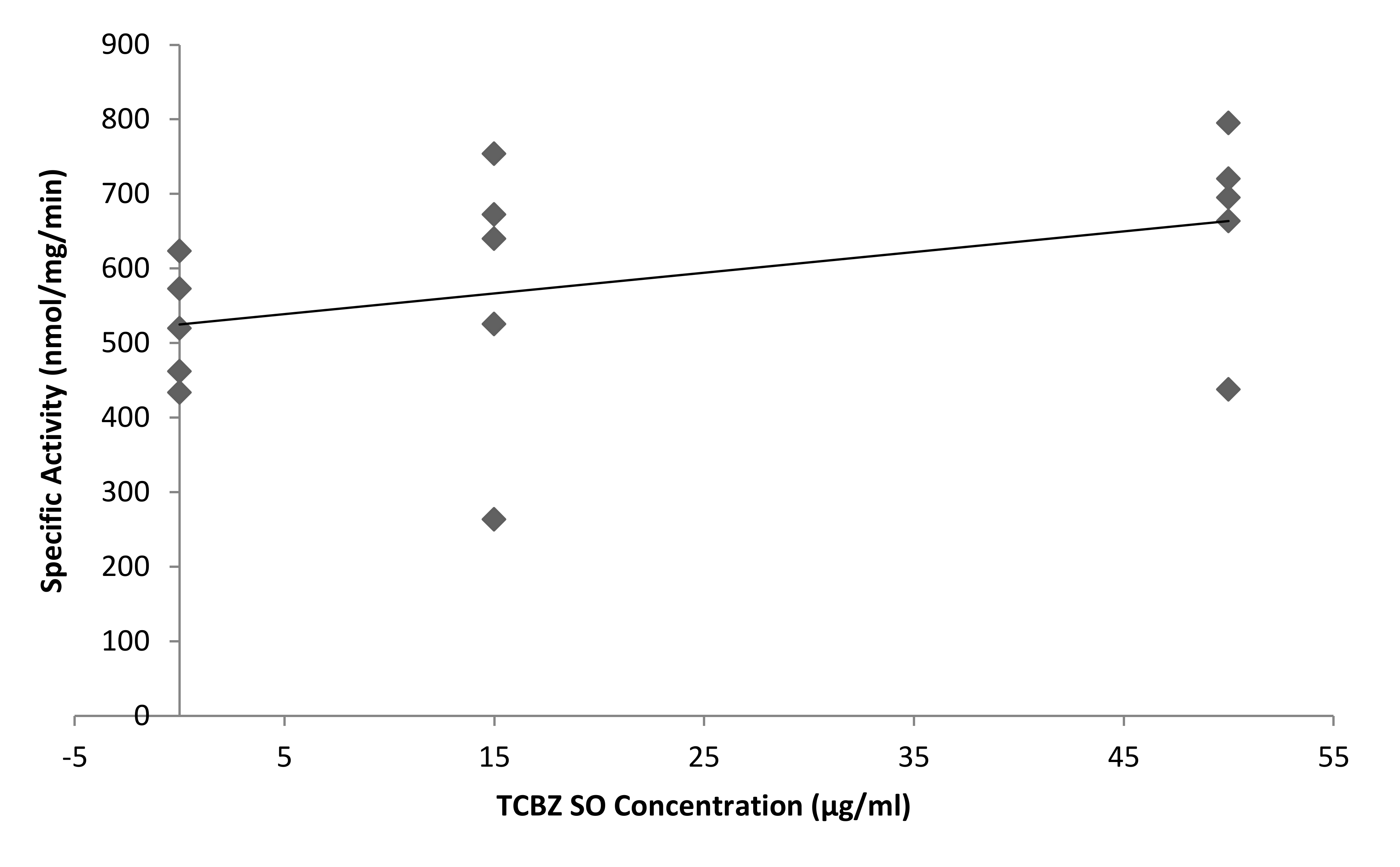

### Supplemental Figure 2

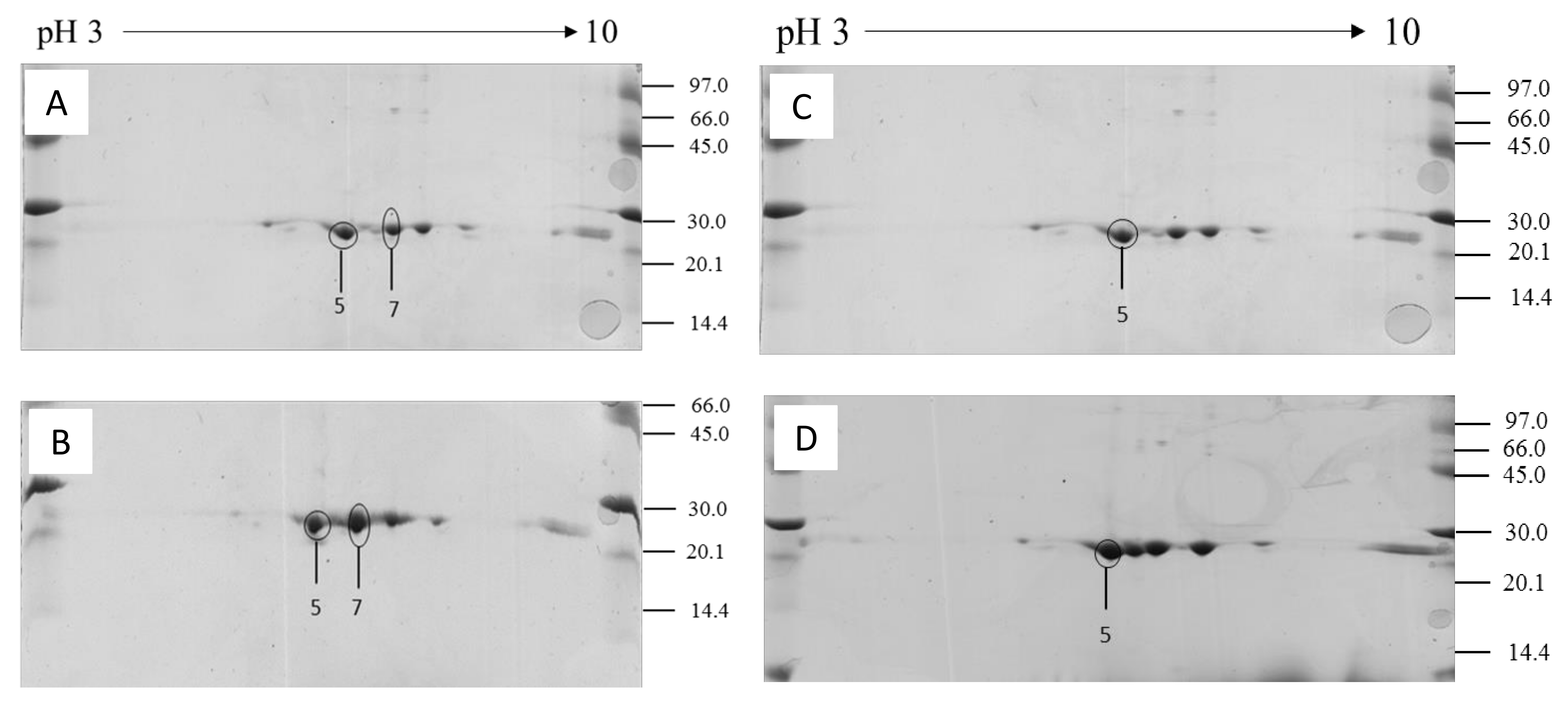

### Supplemental Figure 3

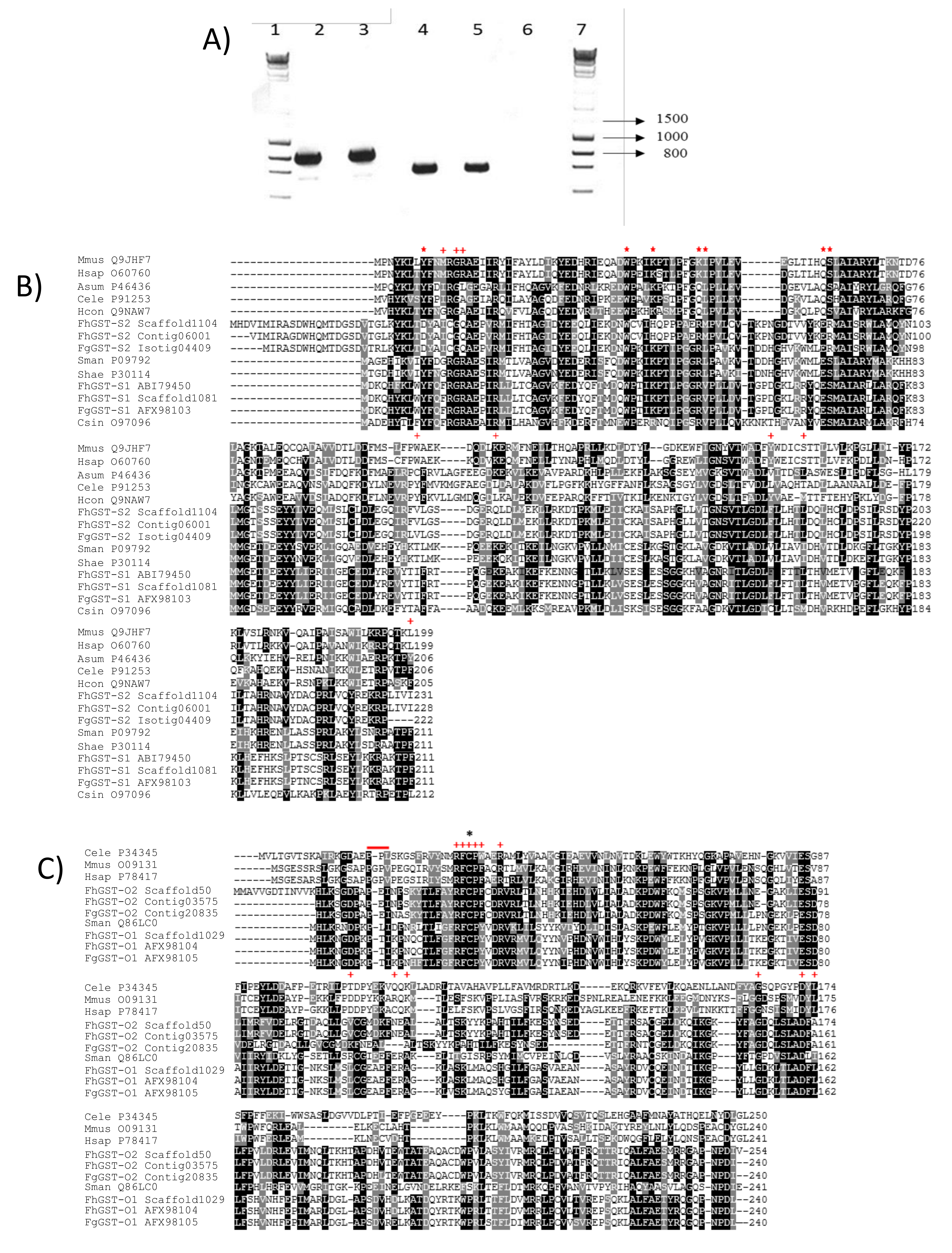

### Supplemental Figure 4

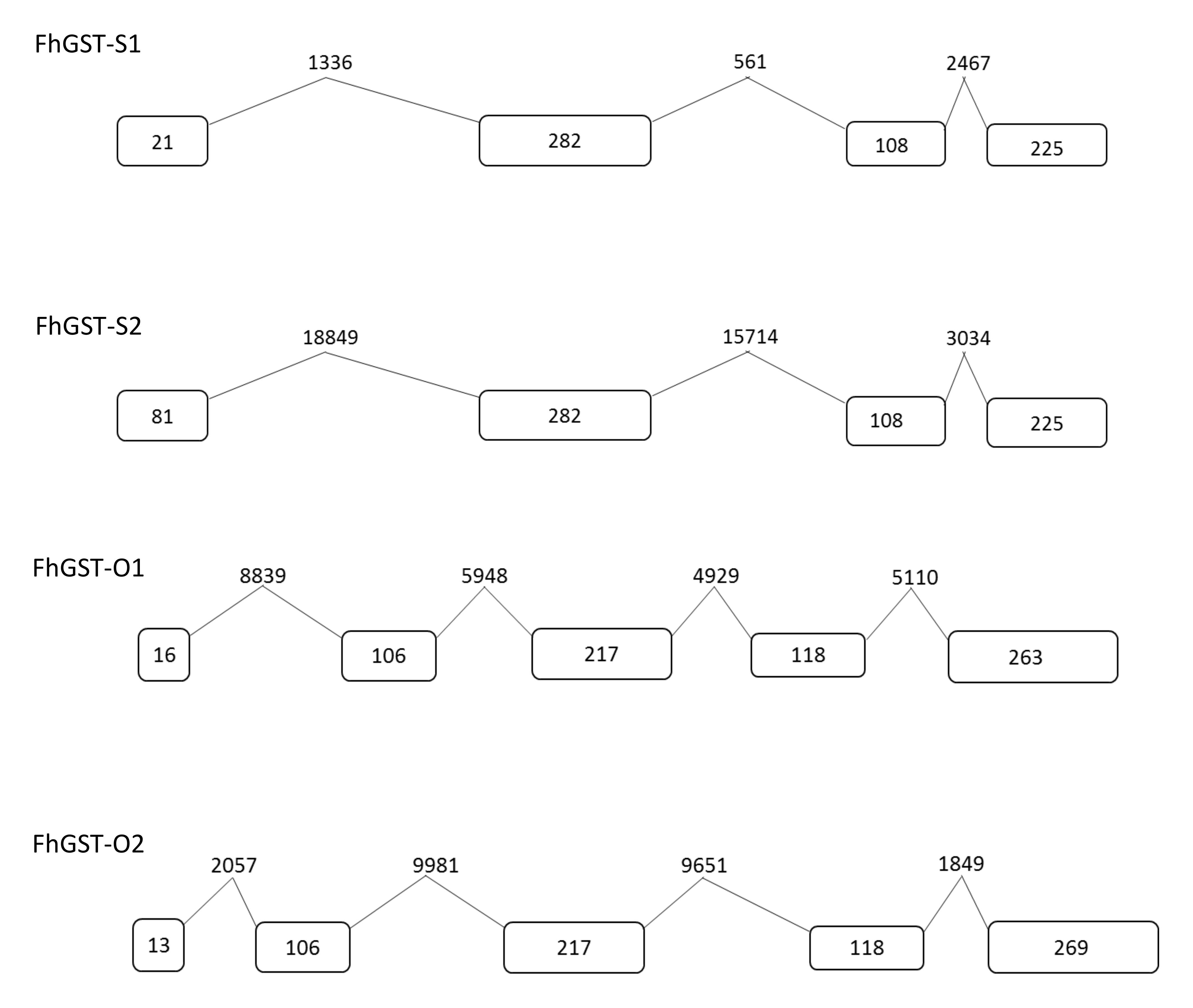
